# Optimizing Artificial Neural Network Models to Predict Brain-Age from Functional MRI

**DOI:** 10.1101/2025.03.28.645964

**Authors:** Mattson Ogg, Lindsey Kitchell

## Abstract

Large MRI datasets combined with deep learning methods have realized a new state of the art for brain-age prediction. Age prediction may serve as a valuable biomarker for brain health and disease given that over-estimated age based on MRI (usually as predicted by a machine learning model; sometimes called a “brain-age gap”) has been associated with neurological and psychiatric disorders. However, most of these results have been achieved via the use of high-resolution structural (T1w) MRI scans. Brain-age prediction via deep learning over large volumes of functional MRI (fMRI) data is less well studied, but could help form a bridge between neural health biomarkers observed in MRI and more portable platforms like functional near infrared spectroscopy (fNIRS), which measure a hemodynamic signal similar to fMRI. In this work, we studied how to optimize deep learning model architectures and training pipelines to predict brain-age from resting state fMRI connectivity data. A wide set of pre-processing and model hyperparameters was explored that included varying the number of nodes and the composition of the input functional connectivity matrices, the size, depth and objective functions of the neural network models, and a time series sub-sampling method as a data augmentation strategy. Model performance was evaluated on both an internal validation set of held-out participants (from the multi-study corpus compiled for training), as well as numerous external corpora not seen during training, which comprised healthy controls and clinical participants. Neural network models with a variety of hyperparameter configurations supported accurate brain-age prediction using fMRI and many models generalized effectively to predict the age of healthy individuals among data sets not seen during training (< 8 years mean absolute error on the external validation dataset). However, we report mixed results regarding a brain-age gap for held out clinical populations using these methods, with a gap observed only among neurodegenerative disorders (here, Alzheimer’s disease), and not among psychiatric disorders or patients with traumatic brain injury. This work constitutes a valuable step towards scalable, portable brain-age prediction but highlights a number of areas where additional work and improvements are needed.

## Introduction

Advanced age is a major risk factor for disease across physiological systems (Kaeberlein et al., 2015; Tian et al., 2023) and age-related biological processes have been linked to many prevalent neurological disorders (Hou et al., 2019). Measuring healthy or unhealthy aging trajectories will be critical for assessing interventions designed to slow biological senescence (Kaeberlein et al., 2015) and preserve cognitive function into old age (Carmona & Michan, 2016). The need for these interventions and an improved understanding of the diverse ways that individuals age is acutely important as life spans increase and large portions of the population become at risk for age related diseases (Dorsey et al., 2013; Feigin et al., 2020). Neurological and psychiatric disorders, in particular, are a leading cause of disability that increase in prevalence with age (Feigin et al., 2020; Hou et al., 2019; Van Schependom & D’haeseleer, 2023).

One biomarker that has emerged to quantify age-related brain health is the difference between the observed or predicted age of someone’s brain and their true age, often called a “brain-age gap” (Cole & Franke, 2017; Franke & Gaser, 2019). Advanced biological brain-age (i.e., a larger brain-age gap) has been observed in a number of clinical populations relative to age-matched controls (e.g., Kaufmann et al., 2019, see Franke & Gaser, 2019 for review), often finding that this gap increases alongside more advanced disease or other diagnostic outcomes. The rationale for the efficacy of a brain-age gap biomarker is that many neurological disorders affect the brain via the same mechanisms as normal aging, accelerating an observable set of secondary effects that age estimation models interpret as being older (Cole & Franke, 2017).

Brain-age prediction and brain-age gap biomarkers have been most extensively studied using structural (usually T1-weighted) MRI. Large datasets like the UK Biobank (Miller et al., 2016) allow for training models on tens of thousands of participants and these models support very accurate age estimation, down to around 4 years mean absolute error (MAE; Tian et al., 2023) using features describing brain anatomy such as gray matter volumes and cortical thickness. Pairing these very large datasets with simple 3-dimensional convolutional neural networks that learn features directly from a raw (or nearly raw) T1-weighted image reduces age-prediction error even further to an impressive 2.14 years MAE (Peng et al., 2021). Both approaches have revealed associations between increased brain-age gap derived from age prediction models and neuropsychiatric disorders, particularly Alzheimer’s (AD) and Mild Cognitive Impairment (MCI; Lee et al., 2022; Leonardsen et al., 2022; Tian et al., 2023). Other work predicting brain-age using models trained on large structural MRI datasets has shown an association between increased brain-age gap and schizophrenia, multiple sclerosis, and bipolar disorder (Kaufmann et al., 2019), as well as traumatic brain injury (TBI; Cole et al., 2015; Spitz et al., 2022).

When interpreting these results, it is important to note that age prediction error typically increases by at least a few years when models are tested on new datasets comprising new scanners or participant populations not seen in training (e.g., Leonardsen et al., 2022). (Dufumier and colleagues (2022) describe this issue in great detail and suggest the use of two validation stages. The first is described as an *internal* validation sample, made up of subjects not seen by a model during training, but whose scans were obtained from the same population sample or imaging center as the training data (essentially a held-out split of the training data). A second *external* validation sample then constitutes a stronger test of model accuracy, and is made up of data from subjects not seen during training, typically data from a completely new neuroimaging dataset or study site. Even for very large datasets, models trained on one or a small number of sites (e.g., the UK Biobank), may contain biases that limit generalizability to new sites or participants. External validation performance can be a valuable method for gauging model generalizability.

Accurate brain-age prediction that relies on MRI also comes with some well-known shortcomings, primarily that the procedure is expensive, contraindicated for some individuals (with implanted metal or claustrophobia), and is not portable. Some work has been done to derive brain-age estimates from electrophysiological signals like EEG as a more portable solution. (Engemann and colleagues (2022) compared a variety of models across four proposed benchmark datasets, achieving MAEs on the order of 7-8 years and as low as 6.5 years for some configurations (using cross validation). Sun and colleagues (2019) trained a model using a large corpus of data using features derived from EEG signals obtained while individuals slept and were able to estimate age down to 7.6 years MAE. Participants with neurological or psychiatric conditions were estimated to be around 4 years older than healthy controls in this study, providing evidence for a brain-age gap based on EEG. In more recent work using deep-learning methods (Brink-Kjaer et al., 2022), age estimates based on sleep signals could be reduced to 5.8 years, and were associated with all-cause mortality risk.

Functional near-infrared spectroscopy (fNIRS) is another portable neuroimaging modality. FNIRS measures hemodynamic changes in blood oxygenation rather than directly measuring electrical activity as in EEG (Pinti et al., 2020). Unfortunately, this modality has not been extensively used to investigate brain-age prediction, and it is difficult to compile large databases of fNIRS data from public sources. However, the hemodynamic signal measured by fNIRS is very similar to the BOLD signal measured in fMRI (Schroeter et al., 2006; Steinbrink et al., 2006). Thus, advances in brain-age prediction using fMRI may provide insights that could be leveraged by future, more portable fNIRS-based age prediction applications. This approach has the benefit of allowing machine learning researchers to leverage the large quantities of public fMRI datasets that already exist to support model development and optimization.

Brain-age prediction in fMRI has been more limited than work predicting age from structural MRI, and age prediction errors are higher in fMRI, but similar patterns of a brain-age gap (with more advanced ages predicted among clinical groups) has emerged. Millar and colleagues (2022, 2023) predicted brain-age using fMRI with an accuracy of 8-8.5 years MAE, and increased brain-age gap estimates were associated with worse scores on clinical measures (e.g., Clinical Dementia Rating, or CDR, scales). Gonneaud and colleagues (2021) obtained an age prediction MAE of 11-12 years on their internal and external validation datasets, and observed an age gap associated with Alzheimer’s specific genotypes and mutations. Finally, Tian and colleagues (2023)were able to predict brain-age around 5 years MAE using a large corpus derived from the UK Biobank. Most of these studies used functional brain-network connectivity matrices derived from a parcellated timeseries of a resting state fMRI scan as input to their models. This representation of the fMRI data is useful as it can easily be applied to standard brain parcellation schemes and over scanning protocols that might have different temporal resolutions (i.e., Repetition Times or TRs). However, these studies have primarily not used neural network classifiers (Millar et al., 2022, 2023; Tian et al., 2023), or used feature engineering steps to refine data from the connectivity matrices prior to input to their neural network (Gonneaud et al., 2021). The success of learning age prediction models directly from structural MRI data (Dufumier et al., 2022; Leonardsen et al., 2022; Peng et al., 2021), suggests networks that learn features directly from connectivity data could be a successful, scalable approach, removing the need for additional pre-processing or feature engineering steps. Finally, we note that much of this work on fMRI-based age prediction (with the exception of Gonneaud et al., 2021) was carried out within a given dataset or imaging center (internal validation), so age prediction error could be expected to rise when tested on data from new studies, datasets or imaging centers (external validation).

The goal of the present study was to learn a neural network representation of brain-age directly from input connectivity matrices. A wide parameter sweep was used to optimize network architectures and training pipelines to accurately predict age, and extensive internal and external validation was used to gauge model performance. Based on external validation using healthy controls, the best performing model was used to assess the brain-age gap measure on three data sets with age matched controls, as well as individuals with neuropsychiatric disorders, TBI, and dementia (specifically AD and MCI). We also trained two sets of baseline models to gauge the performance of fMRI-based neural network classifiers. The first baseline (referred to as the “Structural Baseline”), predicted brain-age from the structural MRI scans in our corpora using a recent high performing neural network approach (the Simple Fully Convolutional Network or SFCN; Leonardsen et al., 2022; Peng et al., 2021). The second baseline (referred to as the “Linear Neural Network (NN) Baseline”), predicted brain-age from functional connectivity matrices using a neural network classifier made up of a single linear output layer with no hidden layers that approximated other linear prediction methods in the literature. Since connectivity matrices are already powerful representations for different neuroimaging tasks (Smith et al., 2013), this latter baseline enables the assessment of how useful neural network architectures are for this task above and beyond the input connectivity matrices. The best performing fMRI model was able to predict brain-age among external datasets better than previous methods especially for younger participants, although errors increased particularly among older samples. We also observe a brain-age gap for dementias (specifically AD and MCI), but not for neuropsychiatric disorders or TBI.

## Methods

### Data and pre-processing

We curated a multi-study training dataset from four sources: NKI-Rockland (Nooner et al., 2012, https://fcon_1000.projects.nitrc.org), 1000 Functional Connectomes (Biswal et al., 2010, https://fcon_1000.projects.nitrc.org), OASIS-3 (LaMontagne et al., 2019, https://central.xnat.org), and data from Setton and colleagues (Setton et al., 2023; Spreng et al., 2022; https://openneuro.org/datasets/ds003592). Datasets targeted for use in model training involved large participant samples, data from multiple sites and those comprising a wide age range which contained both resting state fMRI and corresponding structural MRI scans. Participants between 18 and 89 years of age were retained for training. OASIS-3 participants who had a CDR score over 0 at any study visit were excluded to ensure a healthy training sample. Data from the 1000 Functional Connectomes dataset was restricted to 21 sites^1^ with complete, good quality data (some sites contained only structural or functional data or limited brain coverage). If studies were longitudinal, all scans from all visits were included during training (with the subject age at each visit appropriately updated as the training target). A random selection of 440 participants from this training sample was held out as the internal validation dataset.

The external validation dataset targeted a large normative, cross-sectional sample from a comparable age range (The National Institute of Mental Health (NIMH) Intramural Healthy Volunteer Dataset; Nugent et al., 2022, https://openneuro.org/datasets/ds004215). Clinical datasets, used for testing, each comprised healthy participants as well as patients representing neuropsychiatric disorders (UCLA Consortium for Neuropsychiatric Phenomics LA5c Study;Poldrack et al., 2016, https://openneuro.org/datasets/ds000030), traumatic brain injury (TRACK-TBI; Yue et al., 2013, https://fitbir.nih.gov), and dementia (The Alzheimer’s Disease Neuroimaging Initiative (ADNI); Jack et al., 2008, http://adni.loni.usc.edu/) across a wide age range. For the ADNI data the earliest (youngest) scan for each participant was retained. Analyses of the TRACK-TBI were limited to each participant’s first scanning run (although using the second obtained the same results). Control participants in the LA5c and ADNI cohorts were slightly younger than the clinical cases. Thus, for these datasets only half the controls below the median age were retained. This achieved evaluation sets with no significant age differences between control and clinical samples (all *t*-test and Wilcoxon tests *n.s.*, *p* > 0.1; for the ADNI data, half the ‘MCI’ labeled participants with ages below the median were also subsequently removed to match the control sample). Demographic descriptors of the final datasets used for model evaluation are presented in Table 1. For all sets of data, only resting state fMRI scans were used, ignoring any task runs.

**Table 1.**

|  | <u>Training</u> |  |  |  |  | <u>Validation</u> |  | <u>Test</u> |  |  |
| --- | --- | --- | --- | --- | --- | --- | --- | --- | --- | --- |
|  | Overall | NKI-Rockland | 1000<br>Functional<br>Connectomes | OASIS-3 | ds003592 | Internal<br>Held-out Train | External<br>NIMH Healthy | LA5c | TRACK-TBI | ADNI |
| Age (Mean/Median) | 49/54 | 49/52 | 25/22 | 67/68 | 40/26 | 46/48 | 34/30 | 35/33 | 39/36 | 74/75 |
| Age Range (Min./Max.) | 18/89 | 18/85 | 18/65 | 42/89 | 18/89 | 18/88 | 20/72 | 21/50 | 18/71 | 55/89 |
| Gender (M/F) | 797/1,363 | 268/556 | 246/366 | 189/304 | 94/137 | 208/232 | 46/84 | 134/96 | 334/172 | 375/386 |
| Unique Participants | 2,161 | 825 | 612 | 493 | 231 | 440 | 130 | 230 | 506 | 761 |
| Number Functional Scanning Runs | 6,387 | 3,490 | 612 | 1,824 | 461 | 870 | 130 | 230 | 506 | 761 |
| Total Training Examples (with Subsampling) | 54,161 | 31,356 | 4,520 | 12,753 | 5,532 | N/A | N/A | N/A | N/A | N/A |

All data were minimally pre-processed using fmriprep (version 20.2.6; Esteban et al., 2019) to obtain a version of each functional run in MNI152 space as well as a corresponding set of confounds. We executed fmriprep with flags to ignore slice timing, disable freesurfer and to use susceptibility distortion correction. We provide fmriprep boilerplate language representative of each dataset in the supplemental material. Further pre-processing was carried out using the nilearn package (version 0.8.1) in python. Each functional run processed by fmriprep was parcellated using the 100, 200 and 300-node Schaefer atlases (Schaefer et al., 2018 similar to the 17-network atlas, Yeo et al., 2011). Data were then standardized (z-scored), detrended and lowpass filtered at 0.08 Hz. The confounds file output by fmriprep was used to regress out 12 motion parameters, csf, white matter and global signal. The first 5 samples from each functional timeseries were then removed to avoid any scanner stabilization artifacts. From these functional timeseries we calculated connectivity matrices composed of either standard or partial correlations. These connectivity matrices were used as input for the neural network models as well as the Linear NN Baseline. A temporal sub-sampling augmentation was implemented by deriving connectivity matrices in 100-second windows, sliding the window through the timeseries 50-seconds at a time. Data from any scans that contained fewer than 100 TRs were excluded. If a functional run was not successfully processed by fmriprep or did not result in a confounds file it was ignored.

For the data used in the Structural Baseline model, the anatomical scans above (corresponding to the functional runs) were processed through fmriprep. If a participant only had one study visit (i.e., one session where they were the same age), then the original anatomical outputs from fmriprep (from above) were used. For datasets that involved more than one study visit (i.e., where multiple structural scans might be present, potentially across years), structural data were re-run at the individual session level with additional flags to process only the anatomical data and to disable freesurfer. This resulted in a minimally pre-processed T1 scan from each session aligned to MNI152 space. We then masked the output from fmriprep, tightly cropped the image and then spatially resampled to a dimensionality of 160x192x160 using linear interpolation (similar to Peng et al., 2021).

### Age model training

#### Functional MRI Model Architecture Search

Input to the functional MRI models of brain age were the connectivity matrices derived from the fMRI timeseries. We trained a set of ‘shallow’ and ‘deep’ feedforward neural network models. These models were composed of blocks that contained a linear layer followed by a gelu activation layer and a batch-norm layer (momentum set to 0.1 with no learnable affine parameters). The shallow networks were made up of a single linear block before the output layer. The deep networks contained 2 to 4 linear blocks followed by a final linear block before the output layer. The number of activation units in the deep network layers decreased by half every subsequent linear block, until the final linear block which was the same size as the first layer, thus ensuring the dimensions of the penultimate embeddings (i.e., the layer just before the output layer) of both the shallow and deep networks were consistent. All linear layers were initialized with a Kaiming normal function. All models were trained for 30 epochs with a batch size of 16, a learning rate of 0.001, and an Adam optimizer with a weight decay of 0.0001. A learning rate scheduler was used that reduced the learning rate if no validation loss improvement was observed for three epochs. All neural network training was performed using pytorch (version 1.9.1) and pytorch-lightning (version 1.5.10). An example of a configuration of the fMRI model training pipeline is provided in Figure 1.

**Figure 1.**
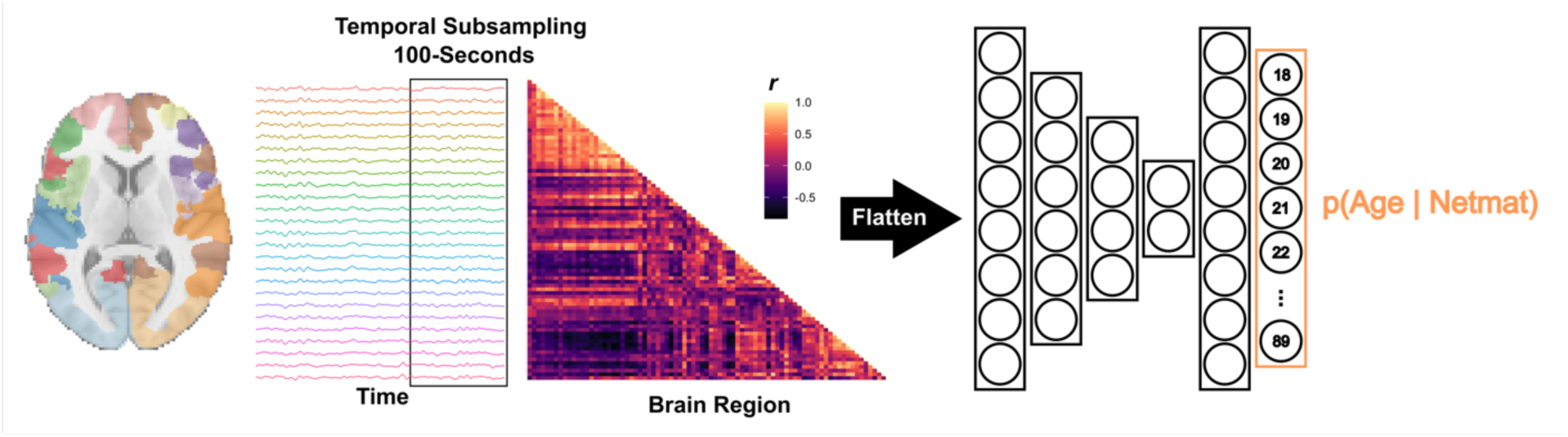
Schematic of an example ‘deep,’ 4-layer feedforward age-prediction model pipeline which uses a classification objective function and takes a 300-node fully connected network connectivity matrix as its input.

Classification and regression-based variants of the age-prediction models were explored via the use of different output layers and objective functions. The classification models contained an output unit for each integer (age) between 18 and 89. These were trained to minimize cross entropy loss for predicting the age of the participants at each scan (rounded to the nearest integer). A class-wise (i.e., age-wise) weighting was applied when calculating the cross-entropy loss to account for imbalances across the age distribution. Regression models used a single output unit and were trained with mean-squared error loss to predict age continuously. Scaling (dividing) the predicted and actual age values by 100 when calculating the mean-squared error loss was implemented as it helped stabilize training.

Training runs swept over a number of training configurations and model hyperparameters. Model depth (shallow, then deep models with 2-4 layers) and model size (first/embedding layers of 4096, 2048, 1024, 512, 256, 128, or 64 units) were both varied over different training runs. Changes in model structure were crossed with whether the input connectivity matrices were composed of standard or partial Pearson correlations and 100, 200 or 300 atlas parcellation nodes. Finally, the effectiveness of temporal sub-sampling was explored along with whether a regression or classification loss was applied for age-prediction training.

#### Linear NN Baseline Model

Given that the functional connectivity matrices these neural networks used as input are already powerful representations of fMRI data, we wanted to implement a baseline model that could help determine what the neural-network architectures contributed to age prediction over and above these functional connectivity matrices. To do this, a set of fully connected functional baseline models was trained that comprised the same training routines as the classification and regression models described above, only these baseline models excluded the model depth and model size conditions because they contained no hidden layers. That is, they were trained to linearly map the contents of each functional connectivity matrix to the corresponding output units.

#### Functional MRI Model Validation

Models were validated in two phases. For each set of hyperparameters, the model weights from the epoch that achieved the lowest mean absolute error on the internal validation participants (i.e., those held out from the training corpora) was retained. This best model epoch was evaluated using the external validation dataset (NIMH Healthy Volunteers). The model that achieved the lowest absolute error on these external data was used for the clinical data and brain-age gap analyses. Note connectivity matrices derived from the full duration scan time series (i.e., no temporally subsampled scans) was used for all validation and testing data.

#### Functional MRI Model Ablation Analyses

Finally, ablation analyses were conducted to assess what functional brain networks were most important for predicting brain-age using the model that best predicted age on the external validation data. For this, the information from the input connectivity matrices corresponding to one of the 17 Yeo networks (i.e., the rows and columns corresponding to the nodes within each network) was removed (zero-ed out). After a network from the input connectivity matrix was removed, the model was re-assessed on the external validation data, and changes in mean absolute error on that dataset from performance on the unperturbed data were recorded. This process iterated through the 17 Yeo networks one at a time. The networks that, when removed, incur the largest increase in error, were interpreted as being most useful for the model when determining a participant’s age (see Lee et al., 2022 for a similar procedure).

#### Structural Baseline Model

The SFCN model (See Leonardsen et al., 2022; Peng et al., 2021 for more details) was used to compare functional age prediction performance with a more widely studied age prediction approach based on structural T1-weighted scans. Initial experiments using public checkpoints^2^ did not produce good results on our data likely due to differences in pre-processing. However, given that we had a substantial amount of training data we trained a new SFCN model from scratch, which produced much better performance. This new model also provides a more direct comparison of T1w and fMRI age prediction performance since the same corpora were used for model training and evaluation. Our specific SFCN implementation comprised six three-dimensional convolution blocks made up of a three-dimensional convolution (kernel size = 3, stride = 1, padding =1), batch-norm (momentum = 0.1) and max pooling (kernel size = 2, stride = 2, dilation = 0, padding = 1) followed by a gelu activation layer. These layers contained 32, 64, 128, 256, 256, and 64 channels. The last block had no max pooling. The output of the convolutional blocks was then average-pooled followed by 50% drop-out and a final convolutional layer connected to a single output unit. This model was trained in a regression set up to minimize mean-squared error (with scaling as above) for 80 epochs at a learning rate of 0.01, and a batch size of 8 using the same Adam optimizer and learning rate scheduler as above.

### Brain-age gap assessment

To evaluate a brain-age gap biomarker, the best performing model was used to evaluate the held-out clinical datasets LA5c, TRACK-TBI and ADNI (not used during training). These clinical datasets were curated such that there was no significant difference in age between the healthy controls and clinical samples. After evaluation, the model predictions were subtracted from the participants’ actual ages to obtain a brain-age gap (sometimes called brain-age delta). Following Millar and colleagues (2022, 2023), brain-age gap was assessed within a linear regression model, which residualized brain-age gap relative to chronological age, and to statistically control for other variables that might introduce bias. Specifically, this linear model predicts brain-age gap as a function of a variable coding for a participants’ clinical state (or a clinical outcome scale) as well as a set of variables of no interest included to offset any potential biases: chronological age (to residualize well known age-wise biases in brain-age prediction; see Smith et al., 2019 for discussion), gender, and study-site (where available). The study-site predictor is coded as a factor with the site with the largest number of participants included as the intercept. Predictors for clinical state are factors with levels corresponding to each clinical group and the healthy control group is always coded as the intercept.

LA5c comprises one site (UCLA), so this variable is not included, and we examined a predictor for clinical cohorts (Control, Schizophrenia, ADHD, Bipolar). For TRACK-TBI we examined clinical group variables (Control vs TBI) as well as outcome measures like Glasgow Coma Scale (GCS), Glasgow Outcome Scale-Extended (GOSE) and Disability Rating Scale (DRS), each within a separate linear model. For ADNI we assessed clinical groups (Control, Subjective Memory Concerns (SMC), Early MCI, MCI, Late MCI, and Alzheimer’s disease (AD)) as well as Mini Mental State Exam (MMSE), Montreal Cognitive Assessment (MOCA) and Clinical Dementia Rating (CDR) scales, along with CDR-Sum of Boxes (CDR-SB, which may provide better differentiation among dementia sub-classes than the full CDR score; O’Bryant et al., 2013; Tzeng et al., 2022), again each within a separate linear model. Model beta values allow us to interpret associations between increased brain-age gap and different clinical sub-classes or scales after controlling for variables on no-interest.

## Results

### Model training and validation

Age-prediction models were trained on a large corpus of over 6,000 functional scans from over 2,000 healthy individuals compiled from publicly available datasets (see Table 1). Data from an additional 440 participants from these datasets was held out as an internal validation set to monitor training. A series of different training runs compared a large number of parameters relevant to the model input (composition of the functional connectivity matrices) and artificial neural network model architecture (model size and training objectives). The accuracy of each model on the internal validation dataset as a function of training epoch for all training runs is displayed in Figure 2.

**Figure 2.**
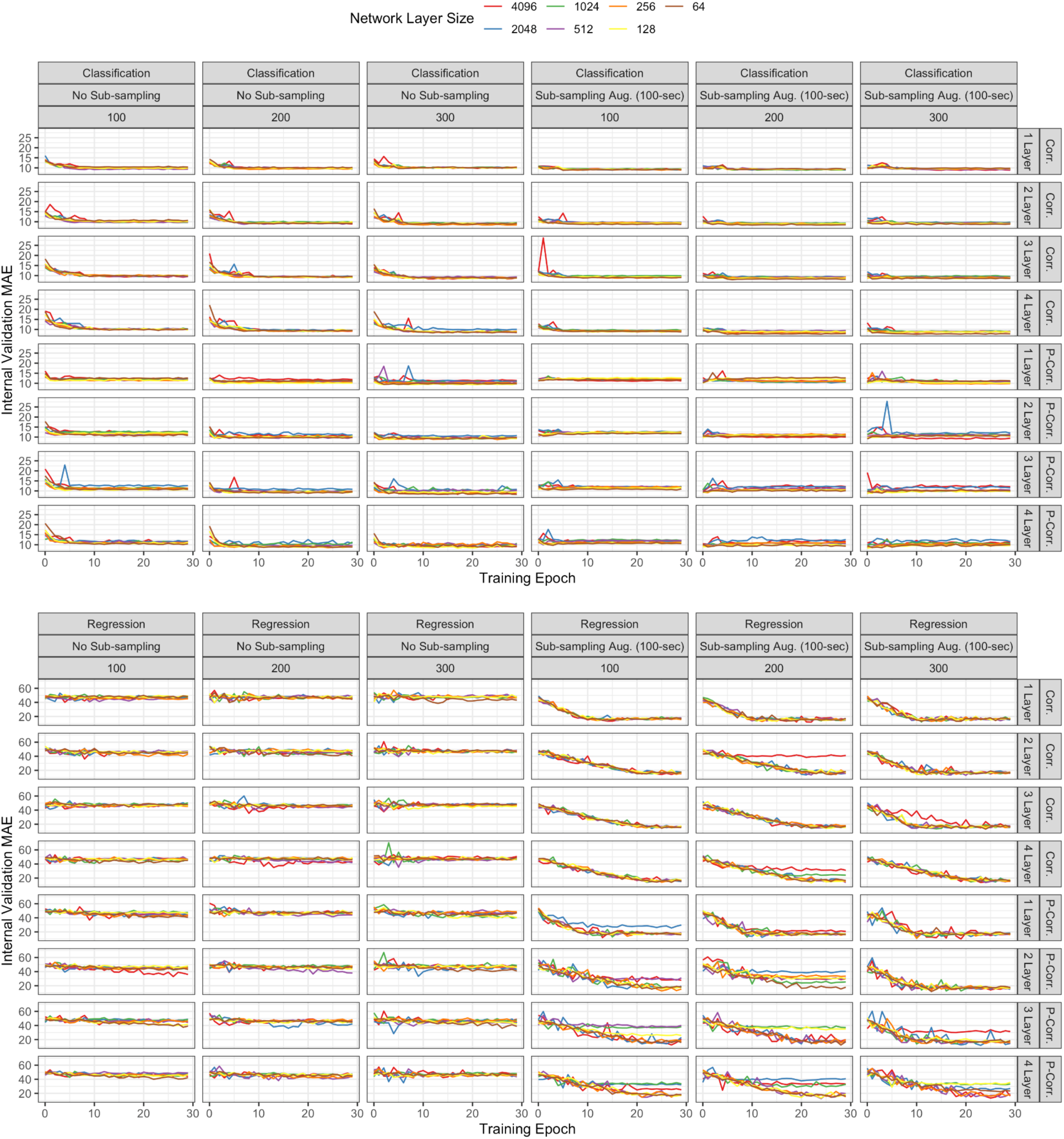
Brain-age prediction results on internal validation data across training epochs. Top figure shows classification model results, while the bottom figure reports results of training runs using regression models and objectives (note the different scale of the y-axes). Columns in both panels are broadly organized by the presence or absence of temporal sampling and then by the number of nodes in the input brain parcellation. Rows in both panels are organized by whether the input network connectivity matrices were populated by a Pearson correlation or a Partial Pearson correlation between node, and the depth of the age-prediction network. Lines are colored to indicate the size of the age-prediction network.

Classification models perform well on the internal validation data across most training configurations and hyperparameters (with or without temporal subsampling), achieving lower than 10 years MAE in many cases. However, temporal subsampling appears to be critical for good regression model performance, and internal validation error is typically higher among regression models than for classification models with otherwise similar hyperparameters. Note however, that the classification models have more parameters than their regression-based counterparts, since the fully connected layer between the penultimate model embedding and the output unit for *each* age (compared to a single regression output unit) increases model size, which may lead to overfitting^3^. Classification models also may not explicitly learn a high level, rank or order-aware representation of age since the output units and cross-entropy objective have no explicit constraints regarding order among adjacent age-wise output units outside the explicit training labels, which could make their learned representations brittle. Regression models explicitly model ordinal relationships among ages, with fewer parameters (a single output unit). In any case, a more critical test of model performance is the ability to generalize to new, unseen data.

Model selection and brain-age gap analyses were carried out in light of known generalization and robustness pitfalls where age prediction errors increase on external data (e.g., Dufumier et al., 2022). To determine which models generalized best to data not seen during training, a second stage of validation was undertaken using an external dataset comprising participants with a similar age range using The National Institute of Mental Health (NIMH) Intramural Healthy Volunteer Dataset (Nugent et al., 2022). For each model and training configuration the weights corresponding to the epoch that achieved the best performance on the internal validation data (i.e., lowest mean absolute error) was retained, and each of these models was evaluated using the external validation data from the NIMH corpus.

These results demonstrated that regression-based models generalized more accurately than classification models. The best performing model (the one that achieved the lowest mean absolute error) on the external validation data used a regression architecture and training objective with temporally subsampled examples during training. The input connectivity matrix was based on the 300-node parcellation with *partial* correlations between each node. The model was three linear blocks deep with a 1024-unit input and embedding layers. Internal and external validation performance for this model is shown in Figure 3. This model achieved an MAE of 7.29 (*r* = 0.64, *p* < 0.001, Bonferroni-corrected for 674 models tested) on the external validation data, but only 13.86 MAE (*r* = 0.79, Bonferroni-corrected *p* < 0.001) on the internal validation data. This model (and indeed all models tested) exhibited the well-known mean-regression bias common to brain-age prediction outputs where the age of younger participants is overestimated and the age of older participants is underestimated (see Smith et al., 2019 for discussion). Interestingly, and by way of comparison, the classification model that performed best on the external validation data comprised the same large, 300-node input connectivity matrix features but processed this input through a very small network, using only 64-unit input and embedding layers, a 4-layer architecture and no temporal subsampling, suggesting that a large amount of regularization (via a small model) helps classification models perform better on external data. This model achieved low internal validation error (8.74 MAE, *r* = 0.84, Bonferroni-corrected *p* < 0.001) but less optimal external validation error (8.14 MAE, *r* = 0.68, Bonferroni-corrected *p* < 0.001). Because all of the datasets with clinical participants that we will use to assess brain-age gap were not seen during training we placed a premium on good external validation performance. Thus, for all subsequent analyses we used the best performing regression model that processes a 300-node partial correlation input, via a 3-layer, 1024-unit feedforward network.

**Figure 3.**
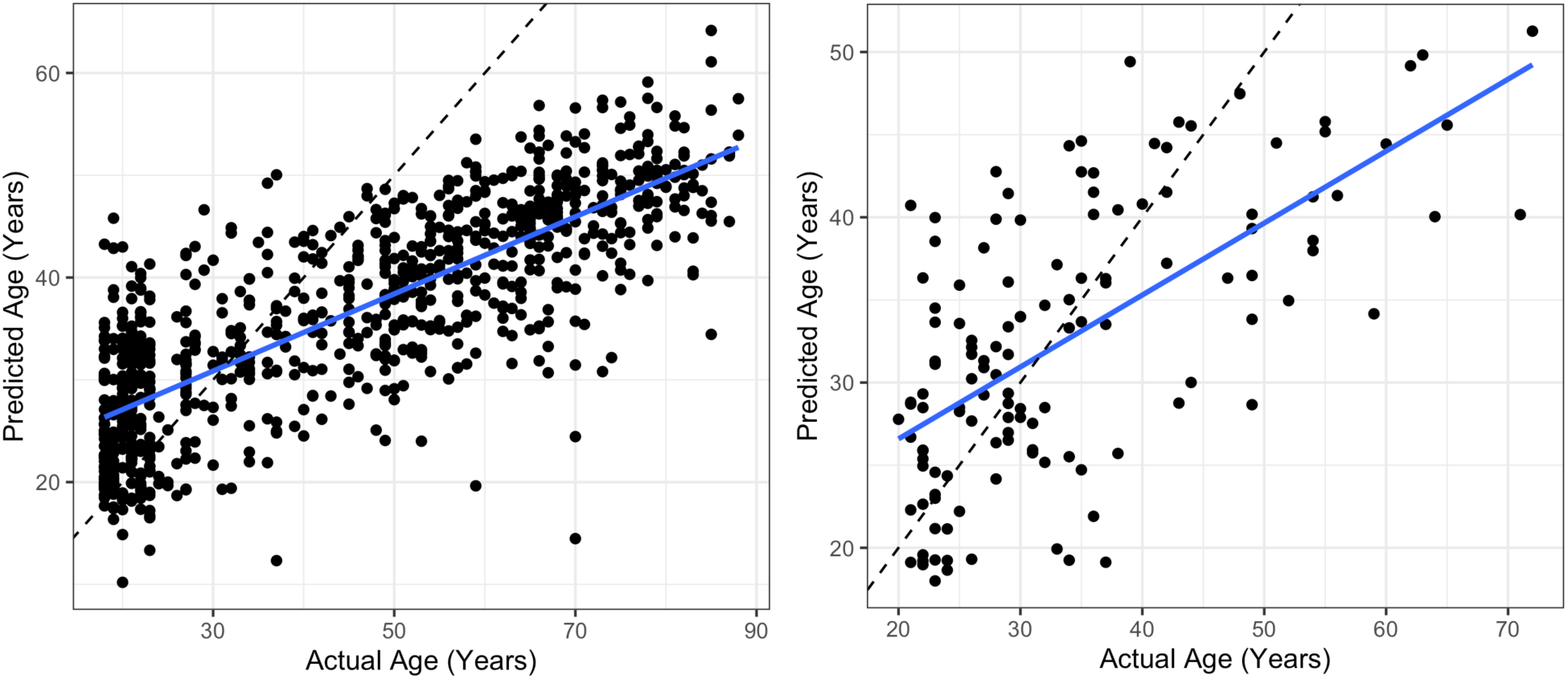
Performance of the best performing brain-age prediction model for healthy participants among the internal validation data (left; MAE = 13.86 years; *r* = 0.79) and the external validation data (right; MAE = 7.29 years; *r* = 0.64). The dotted line in each plot is the identity indicating what would have been perfect performance. Note the different scales of the x and y axes.

Next, we wanted to understand how the best performing fMRI-based age prediction model compared to other age-prediction methods. First, the best functional model was compared to our instantiation of the simple fully connected three-dimensional convolutional network (SFCN), referred to as the Structural Baseline Model, that predicts age based on T1w images. This model was trained from scratch on the T1w images in our training corpus. This model outperformed all functional models on the internal validation data (6.02 MAE, *r* = 0.93, Bonferroni-corrected *p* < 0.001), but slightly underperformed the best functional models on the external validation data (8.79 MAE, *r* = 0.62, Bonferroni-corrected *p* < 0.001). Note, that we expect better performance is possible via a wider search of hyperparameters for the Structural Baseline Model, but since the focus of this report is on functional age prediction models, we relied on this well-studied structural model architecture from the literature (Leonardsen et al., 2022; Peng et al., 2021).

Finally, we wanted to understand whether our deep learning approach offered an improvement over linearly predicting brain-age based on functional connectivity matrices. Functional connectivity matrices were used as deep-learning model inputs, but these are already powerful representations, which have supported accurate brain-age prediction in other work using non-neural network-based modeling approaches (Millar et al., 2022; Tian et al., 2023). Thus, a fully connected baseline model, referred to as the Linear NN Baseline Model, that contained no hidden layers and used the same input connectivity matrix as the best performing model, performed poorly on both the internal validation data (20.36 MAE, *r* = -0.03, Bonferroni-corrected *p* = 1) and the external validation data (26.15 MAE, *r* = -0.20, Bonferroni-corrected *p* = 1). This indicates that the functional model architectures described above were effective for predicting brain-age above and beyond what might be expected based on using the functional connectivity matrices alone. Similarly poor performance was observed across different Linear NN Baseline model hyperparameters (i.e., different input connectivity matrix configurations, and training objectives), Because the fully-connected model performance was very low, we proceed using only our structural, SFCN model as a baseline comparison.

### Localizing networks that influence age prediction

Given good neural-network-based age prediction performance using functional connectivity matrices, we next wanted to understand which brain regions were most important for accurate model predictions. This is partly motivated by the possibility of a future, more portable fNIRS based solution that might have incomplete brain coverage. It would therefore be critical to understand which brain regions are important to target. To do this, the brain regions corresponding to each of the 17-Yeo networks that underpinned the Schaefer atlas were censored and then the external validation data was re-evaluated. The networks that incurred the largest increase in age prediction error (MAE) were interpreted as being most critical for the model when making inferences.

The results of this analysis are presented in Figure 4. Removing regions in the somatomotor networks incurred the largest increase in error (+2.61 years and +1.20 years MAE for SomMotA and SomMotB, respectively), followed by default mode (DefaultA, +0.76 years MAE), dorsal attention (DorsAttnA, +0.71 years MAE) and visual (VisPeri +0.64 years MAE) networks. Previous work by Setton and colleagues (2023) identified changes in these networks as a function of aging. These data were included in our training corpus, so it is likely the model learned similar patterns. However, the Setton data were only a small portion of the training corpus (about 300 out of over 2000 participants). Additionally, this brain-network ablation analysis evaluated performance on a completely separate dataset. Specifically, this functional model was mostly trained on NKI data (since this predominated the training corpus), and ablations to the NIMH data at evaluation time are what produce these patterns. Taken together, our results bolster the previous work by Setton and colleagues (2023) and suggest that age related changes in these networks may be a fairly robust feature of aging in functional connectivity patterns. With the exception of medial portions of motor cortex, most of these networks are located superficially, which bodes well for imaging technologies located on the scalp (such as fNIRS) that may have limited coverage or penetration.

**Figure 4.**
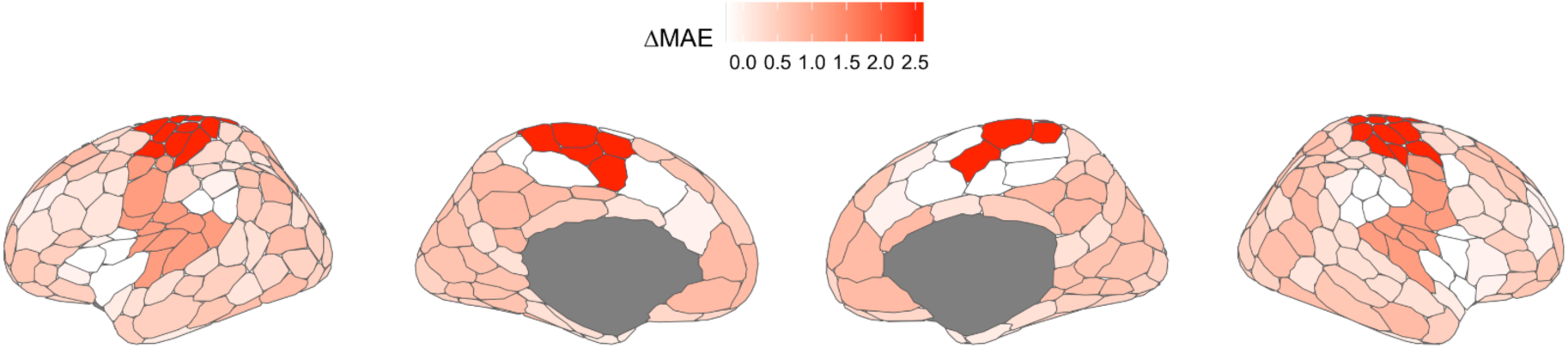
Results of the network ablation studies to determine the most important brain networks and regions for age prediction. Ablations were made network-wise, and the resulting change in MAE is assigned to each of the regions within that network overlaid on the 300-node Schaefer atlas.

### Brain-age gap assessment

We demonstrated that our neural network model can predict the age of individuals from functional connectivity MRI data and that this model can generalize to accurately predict the age of individuals collected via different neuroimaging studies and from different imaging centers. With this robust model in hand, we wanted to investigate a potential clinical biomarker known as the brain-age gap (Cole & Franke, 2017; Franke & Gaser, 2019), where an individual’s biological age is observed (or here, predicted by our model) to be more advanced than the individual’s true, or chronological, age. To assess this biomarker using the functional model’s age estimates, a new collection of datasets (not seen during training or validation) was obtained comprising groups of individuals with a clinical neuropsychiatric diagnosis and healthy, age-matched controls. A brain age gap for clinical patients indicates that they present, based on their functional connectivity data, as being biologically older than they actually are and older than the age-matched cohort of healthy individuals. Rigorous statistical controls helped evaluate the brain-age gap among these groups of clinical participants. Within a linear model, a brain-age gap variable (i.e., the model’s predicted age for each participant minus their actual age) was predicted as a function of variables coding for diagnoses or clinical outcome measures. Each of these linear models also includes the participant’s chronological age in order to residualize the brain-age gap (and to help offset these models’ bias towards the mean in the regression analyses similar to Millar et al., 2022), as well as other potentially confounding variables (gender, study site, etc.), which together provide strong statistical controls for known sources of bias, and together helps more accurately quantify the relationship between brain-age gap and clinical states. Full regression model results are presented in Tables 2 and 3.

**Table 2.** Brain Age Gap Regression Analyses the Best Functional Connectivity Model.

| **LA5c** |  |  |  |  |  |
| --- | --- | --- | --- | --- | --- |
| **Model Predictor** | ***b*** | ***SE*** | ***t*** | ***p*** |  |
| Intercept | 17.64 | 2.18 | 8.09 | < 0.001 | *** |
| Age | -0.54 | 0.06 | -9.42 | < 0.001 | *** |
| Gender | -0.11 | 1.08 | -0.10 | 0.922 |  |
| **ADHD** | -3.64 | 1.51 | -2.42 | 0.016 | * |
| **BIPOLAR** | 1.45 | 1.40 | 1.04 | 0.302 |  |
| **SCHZ** | -1.03 | 1.42 | -0.73 | 0.469 |  |
| **TRACK-TBI** |  |  |  |  |  |
| **Model Predictor** | ***b*** | ***SE*** | ***t*** | ***p*** |  |
| Intercept | 23.12 | 1.23 | 18.73 | < 0.001 | *** |
| Age | -0.67 | 0.02 | -33.28 | < 0.001 | *** |
| Gender | 0.77 | 0.61 | 1.26 | 0.208 |  |
| **TBI Diagnosis** | -0.62 | 0.77 | -0.81 | 0.418 |  |
| Site | ... | ... | ... | ... |  |
| **Model Predictor** | ***b*** | ***SE*** | ***t*** | ***p*** |  |
| Intercept | 19.46 | 3.98 | 4.89 | < 0.001 | *** |
| Age | -0.67 | 0.02 | -33.05 | < 0.001 | *** |
| Gender | 0.77 | 0.61 | 1.26 | 0.209 |  |
| **GCS** | 0.22 | 0.27 | 0.83 | 0.409 |  |
| Site | ... | ... | ... | ... |  |
| **Model Predictor** | ***b*** | ***SE*** | ***t*** | ***p*** |  |
| Intercept | 22.27 | 1.09 | 20.39 | < 0.001 | *** |
| Age | -0.66 | 0.02 | -32.28 | < 0.001 | *** |
| Gender | 0.79 | 0.62 | 1.27 | 0.205 |  |
| **DRS** | 0.29 | 0.18 | 1.65 | 0.101 |  |
| Site | ... | ... | ... | ... |  |
| **Model Predictor** | ***b*** | ***SE*** | ***t*** | ***p*** |  |
| (Intercept) | 23.47 | 3.39 | 6.92 | < 0.001 | *** |
| Age | -0.68 | 0.04 | -19.15 | < 0.001 | *** |
| Gender | 0.97 | 1.12 | 0.87 | 0.385 |  |
| **GOSE** | -0.17 | 0.39 | -0.44 | 0.660 |  |
| Site | ... | ... | ... | ... |  |
| **ADNI** |  |  |  |  |  |
| **Model Predictor** | ***b*** | ***SE*** | ***t*** | ***p*** |  |
| Intercept | 31.17 | 2.14 | 14.54 | < 0.001 | *** |
| Age | -0.79 | 0.03 | -29.26 | < 0.001 | *** |
| Gender | 1.01 | 0.37 | 2.71 | 0.007 | ** |
| **SMC** | 0.13 | 0.71 | 0.18 | 0.855 |  |
| **EMCI** | 1.06 | 0.60 | 1.78 | 0.076 | . |
| **MCI** | 0.53 | 0.56 | 0.94 | 0.347 |  |
| **LMCI** | 0.63 | 0.67 | 0.94 | 0.347 |  |
| **AD** | 1.34 | 0.62 | 2.16 | 0.031 | * |
| Site | ... | ... | ... | ... |  |
| **Model Predictor** | ***b*** | ***SE*** | ***t*** | ***p*** |  |
| Intercept | 35.44 | 2.91 | 12.17 | < 0.001 | *** |
| Age | -0.79 | 0.03 | -29.55 | < 0.001 | *** |
| Gender | 1.06 | 0.37 | 2.86 | 0.004 | ** |
| **MMSE** | -0.12 | 0.07 | -1.69 | 0.092 | . |
| Site | ... | ... | ... | ... |  |
| **Model Predictor** | ***b*** | ***SE*** | ***t*** | ***p*** |  |
| Intercept | 36.20 | 2.38 | 15.24 | < 0.001 | *** |
| Age | -0.80 | 0.03 | -29.44 | < 0.001 | *** |
| Gender | 0.96 | 0.37 | 2.56 | 0.011 | * |
| **MOCA** | -0.16 | 0.04 | -4.06 | < 0.001 | *** |
| Site | ... | ... | ... | ... |  |
| **Model Predictor** | ***b*** | ***SE*** | ***t*** | ***p*** |  |
| Intercept | 31.74 | 4.67 | 6.79 | < 0.001 | *** |
| Age | -0.78 | 0.05 | -14.86 | < 0.001 | *** |
| Gender | 0.99 | 0.66 | 1.51 | 0.133 |  |
| **CDR** | 1.61 | 0.73 | 2.21 | 0.029 | * |
| Site | ... | ... | ... | ... |  |
| **Model Predictor** | ***b*** | ***SE*** | ***t*** | ***p*** |  |
| Intercept | 31.01 | 2.08 | 14.90 | < 0.001 | *** |
| Age | -0.78 | 0.03 | -29.65 | < 0.001 | *** |

**Table 3.**
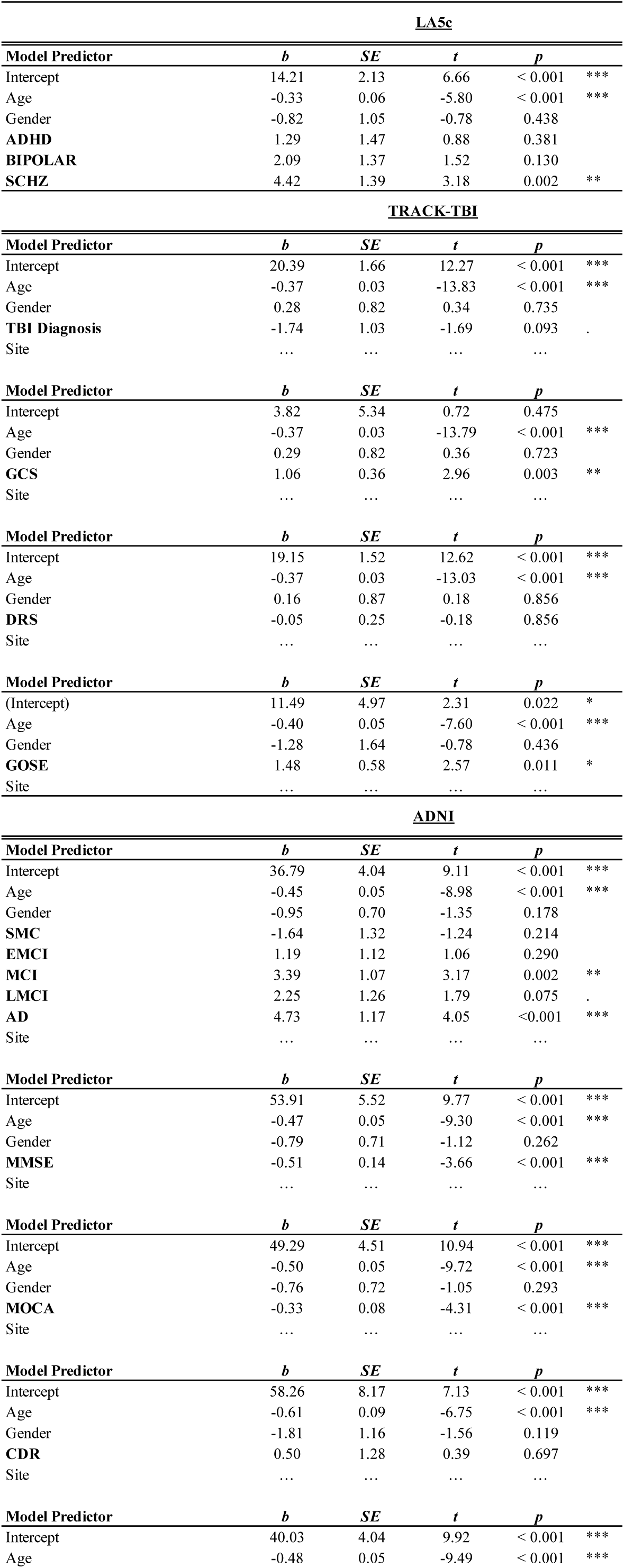
Brain Age Gap Regression Analyses for Structural Baseline Model.

| **LA5c** |  |  |  |  |  |
| --- | --- | --- | --- | --- | --- |
| **Model Predictor** | ***b*** | ***SE*** | ***t*** | ***p*** |  |
| Intercept | 14.21 | 2.13 | 6.66 | < 0.001 | *** |
| Age | -0.33 | 0.06 | -5.80 | < 0.001 | *** |
| Gender | -0.82 | 1.05 | -0.78 | 0.438 |  |
| **ADHD** | 1.29 | 1.47 | 0.88 | 0.381 |  |
| **BIPOLAR** | 2.09 | 1.37 | 1.52 | 0.130 |  |
| **SCHZ** | 4.42 | 1.39 | 3.18 | 0.002 | ** |
| **TRACK-TBI** |  |  |  |  |  |
| **Model Predictor** | ***b*** | ***SE*** | ***t*** | ***p*** |  |
| Intercept | 20.39 | 1.66 | 12.27 | < 0.001 | *** |
| Age | -0.37 | 0.03 | -13.83 | < 0.001 | *** |
| Gender | 0.28 | 0.82 | 0.34 | 0.735 |  |
| **TBI Diagnosis** | -1.74 | 1.03 | -1.69 | 0.093 | . |
| Site | ... | ... | ... | ... |  |
| **Model Predictor** | ***b*** | ***SE*** | ***t*** | ***p*** |  |
| Intercept | 3.82 | 5.34 | 0.72 | 0.475 |  |
| Age | -0.37 | 0.03 | -13.79 | < 0.001 | *** |
| Gender | 0.29 | 0.82 | 0.36 | 0.723 |  |
| **GCS** | 1.06 | 0.36 | 2.96 | 0.003 | ** |
| Site | ... | ... | ... | ... |  |
| **Model Predictor** | ***b*** | ***SE*** | ***t*** | ***p*** |  |
| Intercept | 19.15 | 1.52 | 12.62 | < 0.001 | *** |
| Age | -0.37 | 0.03 | -13.03 | < 0.001 | *** |
| Gender | 0.16 | 0.87 | 0.18 | 0.856 |  |
| **DRS** | -0.05 | 0.25 | -0.18 | 0.856 |  |
| Site | ... | ... | ... | ... |  |
| **Model Predictor** | ***b*** | ***SE*** | ***t*** | ***p*** |  |
| (Intercept) | 11.49 | 4.97 | 2.31 | 0.022 | * |
| Age | -0.40 | 0.05 | -7.60 | < 0.001 | *** |
| Gender | -1.28 | 1.64 | -0.78 | 0.436 |  |
| **GOSE** | 1.48 | 0.58 | 2.57 | 0.011 | * |
| Site | ... | ... | ... | ... |  |
| **ADNI** |  |  |  |  |  |
| **Model Predictor** | ***b*** | ***SE*** | ***t*** | ***p*** |  |
| Intercept | 36.79 | 4.04 | 9.11 | < 0.001 | *** |
| Age | -0.45 | 0.05 | -8.98 | < 0.001 | *** |
| Gender | -0.95 | 0.70 | -1.35 | 0.178 |  |
| **SMC** | -1.64 | 1.32 | -1.24 | 0.214 |  |
| **EMCI** | 1.19 | 1.12 | 1.06 | 0.290 |  |
| **MCI** | 3.39 | 1.07 | 3.17 | 0.002 | ** |
| **LMCI** | 2.25 | 1.26 | 1.79 | 0.075 | . |
| **AD** | 4.73 | 1.17 | 4.05 | <0.001 | *** |
| Site | ... | ... | ... | ... |  |
| **Model Predictor** | ***b*** | ***SE*** | ***t*** | ***p*** |  |
| Intercept | 53.91 | 5.52 | 9.77 | < 0.001 | *** |
| Age | -0.47 | 0.05 | -9.30 | < 0.001 | *** |
| Gender | -0.79 | 0.71 | -1.12 | 0.262 |  |
| **MMSE** | -0.51 | 0.14 | -3.66 | < 0.001 | *** |
| Site | ... | ... | ... | ... |  |
| **Model Predictor** | ***b*** | ***SE*** | ***t*** | ***p*** |  |
| Intercept | 49.29 | 4.51 | 10.94 | < 0.001 | *** |
| Age | -0.50 | 0.05 | -9.72 | < 0.001 | *** |
| Gender | -0.76 | 0.72 | -1.05 | 0.293 |  |
| **MOCA** | -0.33 | 0.08 | -4.31 | < 0.001 | *** |
| Site | ... | ... | ... | ... |  |
| **Model Predictor** | ***b*** | ***SE*** | ***t*** | ***p*** |  |
| Intercept | 58.26 | 8.17 | 7.13 | < 0.001 | *** |
| Age | -0.61 | 0.09 | -6.75 | < 0.001 | *** |
| Gender | -1.81 | 1.16 | -1.56 | 0.119 |  |
| **CDR** | 0.50 | 1.28 | 0.39 | 0.697 |  |
| Site | ... | ... | ... | ... |  |
| **Model Predictor** | ***b*** | ***SE*** | ***t*** | ***p*** |  |
| Intercept | 40.03 | 4.04 | 9.92 | < 0.001 | *** |
| Age | -0.48 | 0.05 | -9.49 | < 0.001 | *** |

#### LA5c

The UCLA Consortium for Neuropsychiatric Phenomics LA5c dataset (Poldrack et al., 2016) is a well-known neuroimaging benchmark dataset containing participants across three psychiatric disorders (schizophrenia, bipolar disorder, and ADHD) as well as healthy controls. Our test set consisted of resting state scans from 230 participants. This test set was curated to achieve a comparable age distribution among control (n=91), schizophrenia (n=50), bipolar (n=49), and ADHD (n=40) groups. Specifically, for these data this meant resolving the slightly lower overall age of the control participants by removing a small number of control participants who fell below the median age of the control group. The linear model used to examine brain-age gap for this dataset contained predictors for chronological age, gender, and diagnosis (no site variable, since all data were from the same UCLA study).

Performance of the best functional model on the LA5c data is shown in Figure 5. This model performed slightly better on healthy controls in the LA5c dataset (MAE = 6.6) than on the NIMH external validation dataset indicating good generalization performance. The model was slightly less accurate for the clinical groups (ADHD MAE = 7.7; Schizophrenia MAE = 9.2; Bipolar MAE

**Figure 5.**
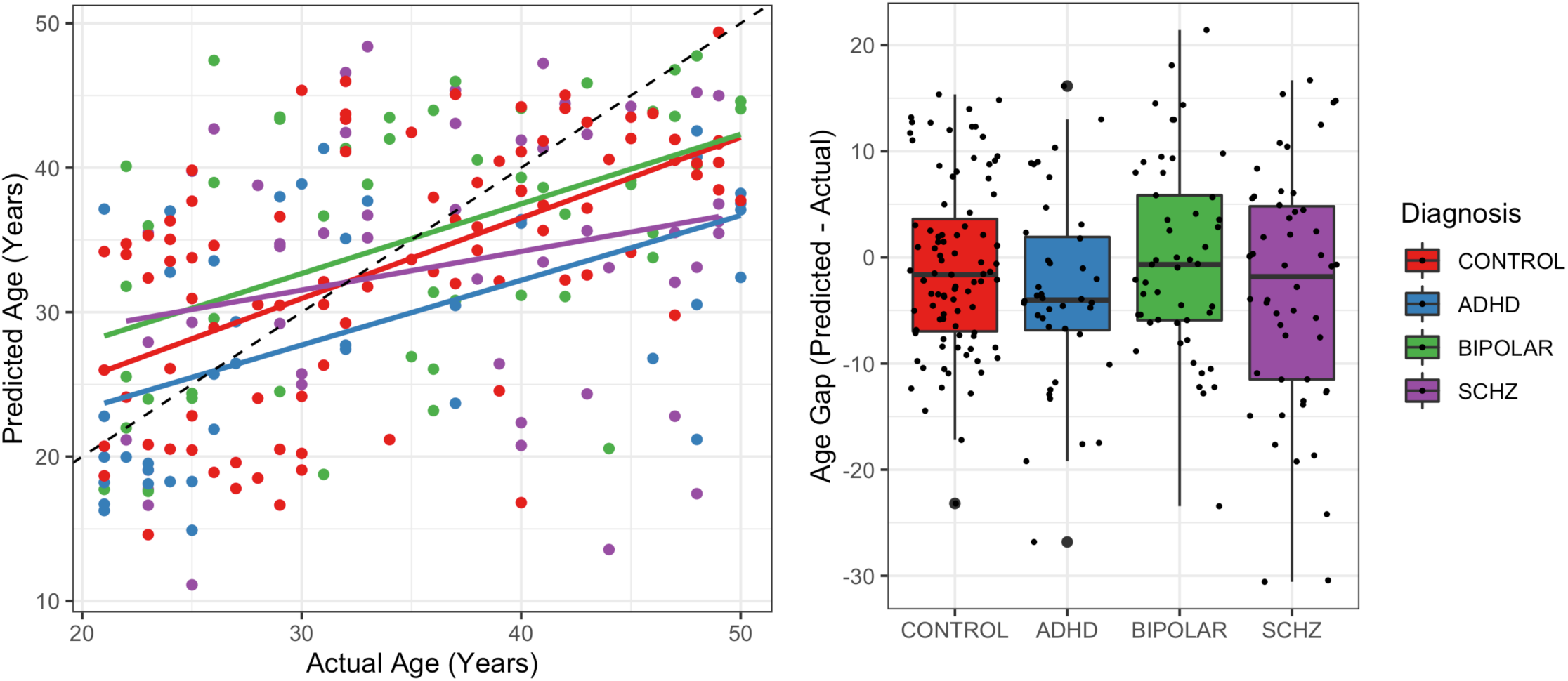
Performance of the best functional age prediction model on the LA5c dataset. Data are colored by each participant’s clinical group. Left: Model predictions plotted as a function of the participant’s chronological age. The dotted line is the identity indicating what would have been perfect performance. Right: Brain-age gap plotted as a function of each clinical cohort.

= 7.3), but the critical test is whether this error is an overestimate (i.e., a brain-age gap) or not. The linear model did not indicate a significant brain-age gap among clinical groups (predictor for bipolar relative to control, *p* > 0.3; predictor for schizophrenia relative to control, *p* > 0.4). In fact, the age of the ADHD group was estimated to be 3.6 years *younger* than the control group (*p* < 0.02).

Finally, the functional age prediction model’s performance on the LA5c data was compared with the Structural Baseline age prediction model. The Structural Baseline age prediction model accurately predicted the age of the control group (MAE = 6.5), and the linear model found a significant brain-age gap where the age of schizophrenia participants was overestimated (relative to controls) by 4.4 years (*p* < 0.002), which aligns with previous findings (Franke & Gaser, 2019; Leonardsen et al., 2022).

#### TRACK-TBI

The TRACK-TBI dataset (Yue et al., 2013) is a large corpus of individuals who have undergone assessment for traumatic brain injury along with healthy controls. This test set comprised 503 participants (all resting state scans, mostly males), 403 of which were diagnosed with TBI and 103 were healthy controls with no significant difference in age between groups. For this dataset, the age gap prediction linear model included chronological age, gender, study site and diagnosis. Follow up analyses substituted the diagnosis variable for Glasgow Coma Scale (GCS), Glasgow Outcome Scale-Extended (GOSE) or Disability Rating Scale (DRS) outcome measures.

Functional model outputs and brain-age gap for the TRACK-TBI dataset are summarized in Figure 6. The functional age prediction model was slightly less accurate than the external validation data in predicting the age of healthy controls (MAE = 10), and accuracy on TBI participants (MAE = 9.8) was comparable. The linear model did not find a significant gap between age predictions for healthy controls and participants with TBI (*p* > 0.4). Follow-up analyses that used continuous clinical outcome predictors (GCS, GOSE, and DRS) instead of the participant’s clinical category also found no relation between clinical outcomes and brain-age gap (all *p* > 0.1). A similar pattern of results was observed using the Structural Baseline model, which achieved slightly worse accuracy than the best functional model on these data, but performance was comparable among healthy control (MAE = 11.9) and TBI (MAE = 11.2) participants. These linear models similarly found no significant difference in brain-age gap between participant groups (*p* > 0.09).

**Figure 6.**
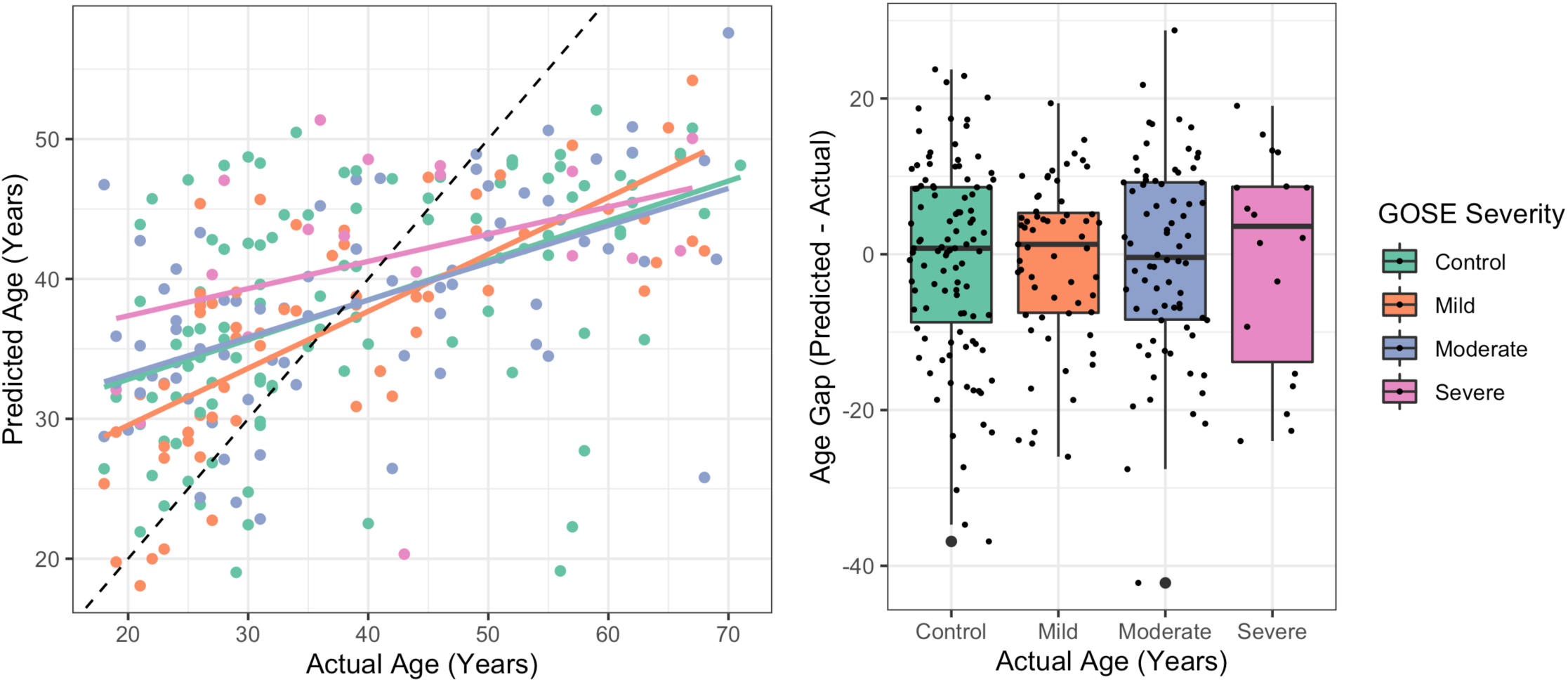
Performance of the best functional age prediction model on the TRACK-TBI dataset. Data are colored by each participant’s clinical group. Left: Model predictions plotted as a function of the participant’s chronological age. The dotted line is the identity indicating what would have been perfect performance. Right: Brain-age gap plotted as a function of each clinical cohort.

#### ADNI

The ADNI dataset (Jack et al., 2008) is a well-studied dataset frequently used to investigate the relationship between brain-age gap and clinical outcomes (e.g.,Gonneaud et al., 2021; Lee et al., 2022; Tian et al., 2023). This test set was made up of resting state scans acquired from 761 participants. Among these data were healthy controls (n=313), patients with subjective memory complaints (SMC; n=65), mild cognitive impairment (MCI; n=126; early, n=98; late, n=71), and Alzheimer’s disease (AD; n=88). Our test set achieved a similar age distribution among these groups. Similar to the LA5c dataset this meant accounting for the lower age of the control participants, again by removing some who were below the median age of this group (and then a similar control among the SMC group to match the new overall age of the controls). Age gap linear modelling analyses for this dataset contained predictors for chronological age, gender, study site and diagnosis. Subsequent analyses replaced the categorial variable of clinical diagnosis group with MMSE, MOCA, CDR and CDR-SB outcome scales.

Functional model performance on the ADNI data along with brain-age gap by clinical group is summarized in Figure 7. The best functional model predicted the age of healthy controls (MAE = 27.8) and clinical participants (MAE = 26.7) worse than previous datasets. However, it can be seen from Figure 7 that this is not entirely unexpected performance given both the advanced age of the entire ADNI cohort (all over 50 years old) relative to our previous data, and the best functional model’s previously discussed bias for underestimating the age of older participants. We note that this is a large degree of error but these biases are partially controlled for in the brain-age gap linear models (where chronological age is included as a predictor of no interest). Additionally, follow-up control experiments were performed to account for this reduced performance among older individuals and the brain-age gap results (see below).

**Figure 7.**
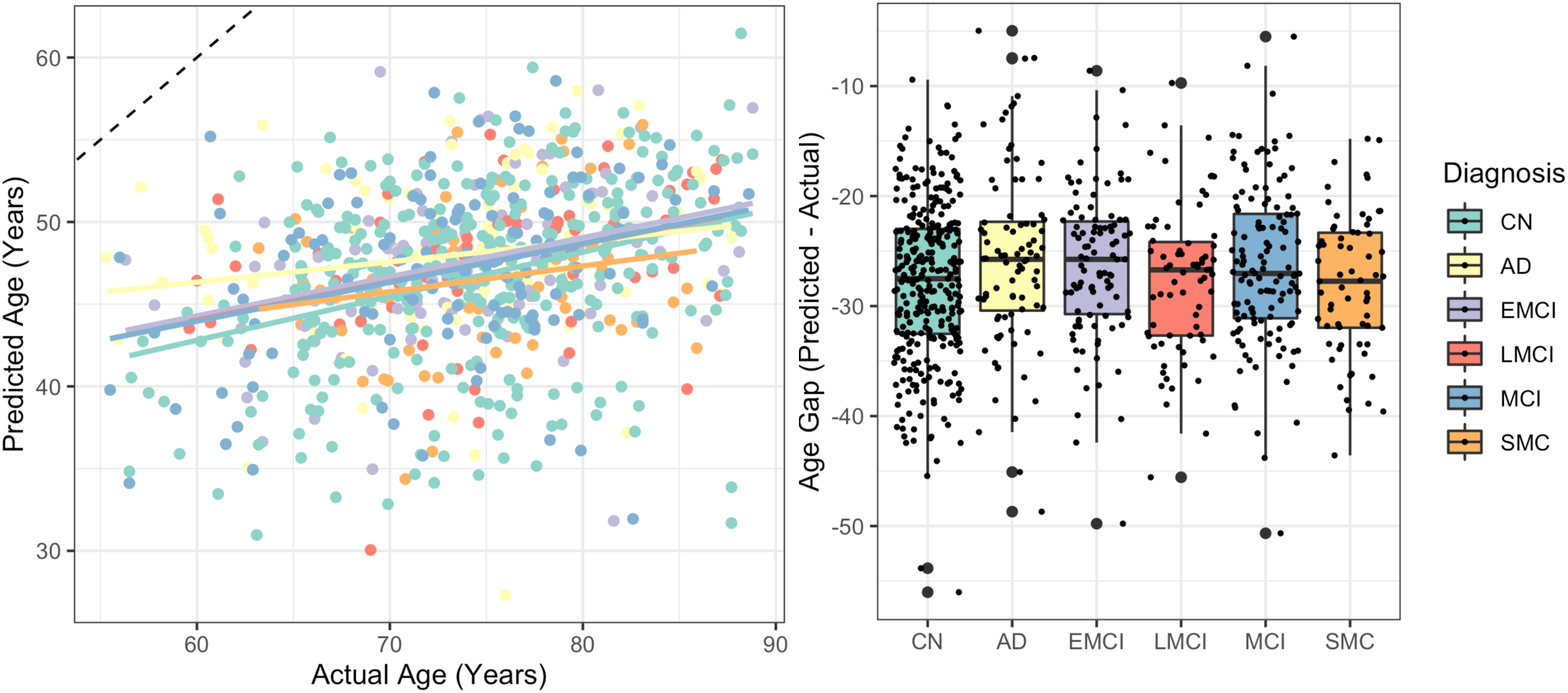
Performance of the best functional age prediction model on the ADNI dataset. Data are colored by each participant’s clinical group. Left: Model predictions plotted as a function of the participant’s chronological age. The dotted line is the identity indicating what would have been perfect performance. Right: Brain-age gap plotted as a function of each clinical cohort.

Linear models revealed a brain-age gap in that the functional age prediction model over-estimated the age of participants with Alzheimer’s disease relative to controls by 1.3 years (*p* < 0.05). Brain-age gap estimates from the functional model also correlated with continuous clinical outcome measures like MOCA scores (-0.16 points on MOCA for every 1 year of brain-age gap, *p* < 0.001) and CDR scores (+1.6 CDR for every 1 year of brain-age gap, *p* < 0.03; +0.28 CDR-SB for every 1 year of brain-age gap, *p* < 0.002), but not MMSE (-0.12 change in MMSE for every 1 year of brain-age gap, *p* > 0.09). In an additional control experiment, a new version of the best performing functional model was trained using only data from older individuals (> 50 years old) in our training corpus (all healthy participants) to account for low performance on this older cohort. This model obtained a similar pattern of results as the original full-age-range model, albeit with slightly reduced error overall.

Finally, we analyzed the output of the Structural Baseline model. This model predicted the age of participants in the ADNI data (control MAE = 8.4; clinical MAE = 9.8) more accurately than the functional model. We also observed a larger brain-age gap of 4.7 years between Alzheimer’s disease participants and controls (*p* < 0.001) and also observed a brain-age gap of 3.4 years between MCI participants and controls (*p* = 0.002). Brain-age gap was also related to clinical outcome measures including MMSE (-0.51 points on MMSE with every 1 year increase in brain-age gap, *p* < 0.001), MOCA (-0.33 points on MOCA with every 1 year increase in brain-age gap, *p* < 0.001), and CDR-SB (+0.44 CDR-SB with every 1 year increase in brain-age gap, p = 0.009; but not full CDR, *p* > 0.6). Similar to the schizophrenia results, these MCI and AD results align with prior work looking at brain-age gap based on structural MRI data (Kaufmann et al., 2019; Leonardsen et al., 2022).

## Discussion

Functional MRI can support accurate brain age prediction, especially when paired with appropriate artificial neural network model architectures. The age of healthy participants from unseen datasets could be predicted to within 7.3 years (and in some cases as low as 6.6 years) and relied on functional activity within the somatomotor, default mode and dorsal attention networks. To our knowledge this is the best reported performance of age prediction using functional MRI on new, unseen datasets. Participants who presented to these models as older were more likely to score worse on clinical outcome measures tracking neurodegenerative diseases (Alzheimer’s and MCI) than age-matched controls, but this brain-age gap effect was not observed for other neuropsychiatric conditions or TBI. In many cases, age prediction performance of the best performing functional model rivaled performance of a state-of-the-art structural MRI modelling architecture trained to predict age on a comparable corpus of healthy participants. However, structural age prediction was more accurate for older participant cohorts (in the ADNI dataset), and overestimation errors from the structural model were associated with schizophrenia diagnoses in addition to a stronger association with clinical measures of Alzheimer’s disease.

Age-estimation using artificial neural network models and fMRI is not well-studied. Thus, a primary goal of this effort was to understand how input and model architecture choices influenced age-prediction errors and generalization. A sweep of parameters related to the construction of the input functional connectivity matrices indicated that a fairly fine 300-node cortical parcellation (which produces large 300-by-300 matrices), provided the best input features for the functional age prediction models. The best-performing regression and classification-based age prediction models also relied on partial correlations among network nodes, rather than standard Pearson correlations. It’s likely that performance could improve further with finer parcellations, but this greatly increases network size (since the input grows in an n-by-n manner) which in turn makes model training more computationally demanding. Input connectivity matrices are extremely powerful, and are currently the inputs to most high performing functional age prediction models (e.g., Millar et al., 2022; Tian et al., 2023) as well as supporting accurate fMRI fingerprinting (Finn et al., 2015; Ogg & Kitchell, 2024; Sarar et al., 2021). However, these results indicate accurate brain-age prediction requires further distillation of these input features. Indeed, models that contained no hidden layers, and simply re-weighted the input connectivity edges did not yield accurate performance. The low performance of the Linear NN Baseline models was surprising since previous work showed these models were a strong baseline for fMRI fingerprinting even up to thousands of participants (Ogg & Kitchell, 2024). In sum, one conclusion from these baseline experiments is that additional layers in the age prediction models allow them to learn more generalized representations of age irrespective of individual variability, that very simple linear models (with no hidden layers) aren’t able to produce.

A sweep of parameters related to the architecture of the age-prediction neural network models compared model depth (which, as discussed above, appears important for age-prediction using functional connectivity matrices) as well as the model outputs and objective functions. Both classification models (where each age rounded to the nearest integer is assigned its own output unit) and regression models (where a single output unit predicts age in a continuous manner) could produce accurate age estimates, but the regression models performed slightly better on external validation data. Because the individual output units of the classification models are fully connected to the embedding layer, these models tended to be much larger than their regression-based counterparts. This increase in free parameters could partially explain our observation that the best classification models performed better on the internal validation data, but performed worse on external validation, indicating a small degree of overfitting. Having discrete output units also requires rounding participant ages, and thus introduces some small error into classification model age estimates. At the same time, cross-entropy objectives for classification models allow for intuitive re-weighting of examples during training that can account for class imbalances, which, to our knowledge, are not available in regression model training regimes. The regression model architectures were overall slightly harder to train, requiring a larger training manifest that included temporally sub-sampled examples to augment the training data in order to obtain good performance. Meanwhile, the top performing classification model was within one MAE of the best regression model on our external validation data, contained many fewer units in each hidden layer and was trained without using temporal sub-sampling. If one is training a model to predict participant ages for a specific site or task, has a limited pre-processing compute budget or does not have to deal with external generalization, classification-based models may offer a good option. However, this is typically not the case, especially as datasets increase in size. Clinical training and evaluation data often require collection via large efforts comprising many study sites. Thus, we placed a premium on good external performance, when validating and selecting the best performing models.

The best performing functional model trained in the course of this work reproduced previously observed brain-age gap findings among Alzheimer’s and dementia patients (Gonneaud et al., 2021; Lee et al., 2022; Millar et al., 2022). However, age-gap effects were not observed for functional models evaluating data from neuropsychiatric patients (schizophrenia, bipolar, ADHD; Kaufmann et al., 2019; Leonardsen et al., 2022) and age-gap effects were not observed for either structural or functional models evaluating data from TBI patients (Cole et al., 2015; Spitz et al., 2022; see Franke & Gaser, 2019 for review). Additional work will be needed to reconcile these results, but we offer a few hypotheses for why these brain-age gap findings differ from prior reports. First, previous work has mostly focused on structural MRI so age-gap effects might be less pronounced in functional connectivity data than structural T1w images. Second, some previous work (but not all) used non-neural network-based age prediction models. Thus, another explanation could be that neural network models may more specifically fit neural aging than the pathology of other conditions, thus failing to produce an age-gap for new data, but still producing reasonable age estimates for healthy controls. A third hypothesis is that this gap could follow from features of our training corpora. We compiled a large multi-study functional imaging dataset for training the age prediction models. However, our set of over 6000 scans from over 2000 participants is still much less than many of the massive datasets used to train structural age prediction models which are on the order of tens of thousands of participants.

This is an ongoing area of discussion that is slightly more advanced in the structural age production literature. For example, standard benchmark corpora now exist for structural age prediction, to help understand site-wise biases in age prediction training (Dufumier et al., 2022), but these do not yet exist for functional age prediction. Absent standard corpora or benchmarks it’s likely research groups will introduce different biases into their model as they compile their individualized training datasets. A final reason that we might have observed only limited a brain-age gap relative to previous work is simply due to the higher error rates we observe with functional models (as low as 6.6 to 7.3 years healthy MAE) relative to structural age prediction models (as low as 2.9 to 3.9 years healthy MAE; Leonardsen et al., 2022). Higher error overall means greater variability in model estimates and more overlap in the hypotheses the model outputs for different clinical cohorts. This makes mean age estimates for a given clinical group harder to separate from the control group.

All datasets used for model training contain some degree of bias and it is worth reflecting on how this might influence the results reported here. First, the functional age prediction model obtained very low errors on some datasets but much higher errors on others. Age-wise biases in age prediction models are well studied (see Smith et al., 2019 for discussion), and we implemented a variety of statistical controls when examining any age-gap produced in the model’s output. However, this range of performance is likely due to the distribution of ages in a given downstream dataset matching the distribution of the training data set (i.e., closer to the mean perhaps like LA5c) or not (as was the case in the older ADNI cohort). Better downstream performance might also be related to how those data do or don’t match other fMRI scanning hyperparameters that are represented in the training data. In some cases, larger age-prediction errors were obtained for some clinical groups relative to controls in the testing datasets (e.g., LA5c), even when there was no brain-age gap observed (i.e., these errors were not systematically overestimations). This could indicate a domain shift between training on healthy controls and evaluating clinical groups, which is a valid and more general criticism of brain-age gap analyses (see Cole & Franke, 2017 for discussion).

Predicting age from functional MRI data using artificial neural networks is still relatively new, and thus offers a number of exciting avenues for future work. First, this report only used very simple neural network model architectures, leveraging a strong set of input connectivity matrices. However, future studies might explore more complex model architectures involving graph neural networks or transformers that could directly model the fMRI timeseries as a sequence learning problem. In an effort to maximize the data used for model training, and reduce potential confounds we focused solely on resting state MRI scans. However, it would be useful to incorporate different tasks in the scanner potentially using a standardized battery designed to maximize age-related differences in neural processing (see Finn, 2021; Finn et al., 2017 for similar discussions related to fMRI connectome fingerprinting). Larger training datasets might also enable improved accuracy (and thus more sensitive brain-age gap detection). Evaluation datasets with more continuous clinical outcome measures (instead of categorical, diagnostic ones) would provide better understanding of the relationship between function brain-age gap and clinical states, support increased statistical power (since the variance is not parcellated by groups), and align with recent pushes towards more continuous definitions of wellness and disease (see Morris et al., 2022). Additionally, longitudinal evaluation datasets could permit an investigation of the stability of age estimates over time.

Finally, this work is a step towards increased portability for age prediction, but these results are still based on hemodynamic signals that require a (non-portable) functional MRI scan. Moving this work towards more portable platforms requires larger corpora of functional near infrared spectroscopy data across the lifespan. These data could help produce methods to adapt or modify functional BOLD hemodynamics recorded in fMRI such that they resemble fNIRS signals. This in turn could support training models on both the large fMRI datasets that are available with the goal of operating on a portable fNIRS platform.

## Conclusion

Brain-age gap may constitute a useful biomarker for assessing neuropsychiatric health and disease. To date, however, most of this research has involved structural MRI, which is bound to large, non-portable imaging devices. In this work, we focused on developing accurate age prediction models using the functional BOLD fMRI signal. We wanted to learn neural network representations for age prediction, directly from input functional connectivity matrices. This functional brain signal is less well studied for age prediction, but may serve as a stepping stone between large scale MRI-based neuroimaging and more portable modalities that measure a similar hemodynamic signal such as fNIRS. We used rigorous internal and external model validation procedures and were able to produce models that can accurately predict brain-age on unseen data better than previous work. In many (but not all) cases our functional models even performed at a similar level as better-studied structural age prediction models trained on comparable data.

## Acknowledgements

The authors acknowledge support from the Independent Research and Development (IRAD) Fund from the Global Health Mission Area of the Johns Hopkins University Applied Physics Laboratory.

## Footnotes

1 ‘AnnArbor_a’, ‘Baltimore’, ‘Bangor’, ‘Beijing_Zang’, ‘Berlin_Margulies’, ‘Cambridge_Buckner’, ‘Leiden_2200’, ‘Milwaukee_a’, ‘Milwaukee_b’, ‘Munchen’, ‘Newark’, ‘NewYork_a’, ‘NewYork_b’, ‘Queensland’, ‘Ontario’, ‘Oulu’, ‘Oxford’, ‘SaintLouis’, ‘PaloAlto’, ‘Taipei_a’, ‘Taipei_b’

2 https://github.com/ha-ha-ha-han/UKBiobank_deep_pretrain

3 Compare: ([embedding-size] * [71-output-units]) and ([embedding-size] * [1-output unit]); for classification and regression models respectively.

